# *De novo* design of selective kinase modulators

**DOI:** 10.64898/2026.07.10.737808

**Authors:** Magnus S Bauer, Saurav Kumar, Mia Donald-Paladino, Dongyang Li, Kody A. Klupt, Matthias Glögl, Ana María Fernández Escamilla, Thomas Schlichthaerle, Edin Muratspahić, Xinru Wang, Ludwig Schmiderer, Sebastian Kenny, Brian Coventry, Bulat Faezov, Wei Chen, Alex F. Shida, Gyu Rie Lee, Yang Hsia, Ryan D. Kibler, Michael B Elowitz, Daniel Lietha, Behnam Nabet, David Baker

## Abstract

Protein kinases are critical regulators of cellular signaling, but precise modulation of their activity remains challenging due to their high structural conservation. Here, we present *de novo* designed genetically encoded miniproteins capable of activating or inhibiting focal adhesion kinase (FAK) by directly targeting the kinase domain itself. Among 96 binders designed to stabilize distinct conformational states of FAK, 33 modulated kinase activity. Biochemical characterization of the four most potent modulators revealed that two designs inhibit FAK with low-nanomolar IC_50_ values while the remaining two potentiated FAK activity by more than two-fold. When expressed in cells, the modulators preserved the same inhibitory and activating effects observed in vitro, establishing that designed conformational binders can directly tune FAK signaling in living cells. Taking advantage of the high similarity between kinases, we redesigned the FAK inhibitors to inhibit Src kinase. Our approach establishes a versatile platform for selective and genetically encoded kinase control as a way to rewire cell signaling and as a starting point for the discovery of novel modulatory sites of kinases.

## Introduction

Protein kinases orchestrate critical signaling processes governing cellular growth, survival, and migration, making them central to numerous diseases, especially cancers (1, 2). Despite their therapeutic relevance, selective targeting of kinases remains difficult due to the highly conserved nature of the ATP-binding domain, often leading to the development of inhibitors with unintended off-target effects against other kinases (3). This promiscuity frequently results in toxicity, thereby limiting therapeutic efficacy. Beyond sequence conservation, kinase catalytic domains are also conformationally plastic: they interconvert among active-like and inactive-like states defined by coupled rearrangements of conserved elements such as the activation segment, including the DFG motif, and the αC helix, which together control productive active-site geometry and catalytic turnover (4–6). Catalysis imposes strong geometric constraints, so active kinase-domain conformations are comparatively conserved, whereas inactive-like conformations are less functionally constrained and therefore structurally heterogeneous (4, 7–9). Consequently, modulators that stabilize specific conformations can, in principle, provide both selectivity, by exploiting state-dependent features beyond the ATP site, and bidirectionality, by shifting pre-existing conformational equilibria toward inhibited or activated kinase-domain states (10). Proteins such as monobodies have been developed to allosterically modulate kinase activity (11), but these methods rely on empirical selection processes and therefore do not provide direct control over where and how a modulator engages the kinase surface.

We reasoned that machine learning-based protein design methods could enable the generation of novel proteins capable of stabilizing defined kinase-domain conformational ensembles. Unlike traditional library screening, computational approaches could allow rapid generation of targeted, optimized binders without extensive experimental iteration. We set out to employ RFdiffusion to generate kinase modulators, selecting focal adhesion kinase (FAK), which is involved in cancer metastasis and cell migration (12, 13), as our initial target. FAK is a good test case for kinase-domain state targeting because available kinase-domain structures show multiple inactive-like geometries rather than a single canonical DFG/αC mode. FAK has not been observed to undergo the αC-helix movement that regulates several other kinase families, and inhibitor-bound FAK kinase-domain structures reveal noncanonical DFG-region conformations, including the helical or semi-flipped DFG conformation induced by TAE226-related inhibitors (14, 15). We aimed to create two classes of binders: one class that would stabilize catalytically impaired, inactive-like kinase-domain ensembles, and another class that would stabilize catalytically competent kinase-domain microstates.

## Results

### Design and testing of *de novo* designed FAK modulators

Based on available FAK kinase-domain crystal structures, we selected an ensemble of conformational templates to sample differences in N-lobe/C-lobe orientation, using ATP-pocket distance as a simple metric for local kinase-domain geometry (Fig. 1ABC). Because the available structures do not fall into a single simple active/inactive classification, we aligned them on the kinase C-lobe and quantified the distance between a glycine-rich-loop reference position in the N-lobe and a DFG/ATP-site-proximal reference point in the C-lobe (Fig. 1B). The phosphorylated kinase-domain structure 2J0L was used as the active-state reference (16), and occupied a compact ATP-pocket geometry, whereas additional kinase-domain structures, including 4D4R, 4D5H, and 4K9Y exhibited progressively larger inter-lobe distances (17, 18). Additional FAK kinase-domain structures, including 1MP8 and 4D55, were included as conformational templates to broaden geometric sampling, but were not treated as formally active-state structures (17, 19). Across this ensemble, we identified solvent-exposed hydrophobic patches near ATP-site-proximal and inter-lobe regulatory surfaces that appeared suitable for *de novo* binder engagement (Supplementary Table T1). We therefore targeted a range of kinase-domain conformations, with the goal of generating binders that could bias FAK catalytic output toward activation or inhibition.

**Figure 1.**
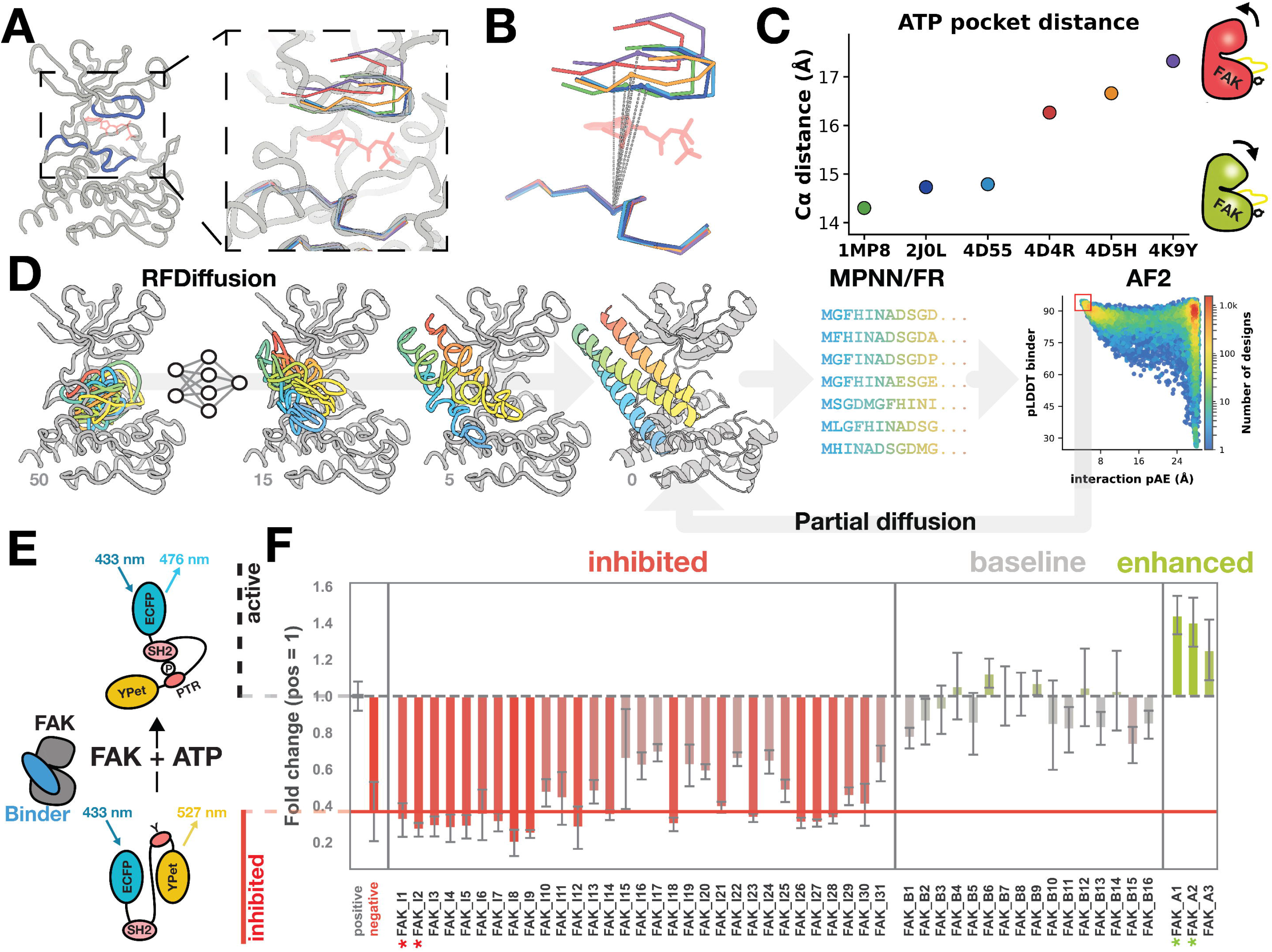
Computational design, structural evaluation, and functional screening of *de novo* FAK modulators. **(A)** Structural comparison of FAK kinase conformations used to guide design. Representative FAK kinase structures are shown after alignment on the kinase C-lobe, with the glycine-rich loop and ATP-site reference elements highlighted and the ATP analog AMP-PNP shown in red. The zoom in view highlights the conformational differences around the ATP-binding pocket relative to the C-lobe. **(B)** Overlay of representative FAK target structures used to quantify ATP-pocket geometry. Structures are aligned on the kinase C-lobe, and the ATP-pocket distance is indicated between residue 430 the glycine-rich-loop reference position and residue 552 in the DFG/ATP-site reference point. AMP-PNP is shown faintly in red to mark the nucleotide-binding site. **(C)** ATP-pocket distance across FAK reference structures. Each point corresponds to one target structure, showing the range of ATP-site-proximal geometries sampled by the FAK kinase-domain templates used for design. The active-state reference 2J0L occupies a relatively compact ATP-pocket geometry similar to 1MP8 and 4D55, while additional templates, including 4D4R, 4D5H, and 4K9Y, span more open geometries. **(D)** Overview of the design pipeline. RFdiffusion generates diverse backbone ensembles which are sequence-designed by ProteinMPNN, relaxed using Rosetta FastRelax and structurally evaluated with AlphaFold2 to select high-confidence FAK binders with low interface pAE and high pLDDT. The best binders underwent another round of partial diffusion followed by ProteinMPNN and scoring with AlphaFold2. An exemplary denoising trajectory (at steps 50, 15, 5, 0) is shown for FAK_I2 in rainbow with the kinase structure (PDB ID: 4D4R) shown in gray. **(E)** Schematic of the FAK FRET biosensor assay. FAK activity is detected by the ECFP-SH2-YPet reporter, which shifts emission depending on substrate phosphorylation. Designed binders that inhibit kinase activity maintain the FRET signal, while binders that favor the active state or remain baseline activity decrease it. **(F)** Primary screen of 50 binder designs for modulatory activity for kinase turn over. Bars show fold change relative to a positive control. The solid red line indicates the inhibited reference and the dashed gray line indicates the activated baseline reference. Four top candidates are highlighted with stars: two inhibitors in red (FAK_I1, FAK_I2) and two activators in green (FAK_A1, FAK_A2), each producing strong and reproducible modulation of FAK activity.

Using RFdiffusion, we generated candidate binder backbone geometries against the selected FAK kinase-domain conformational templates, placing compact scaffolds onto the chosen surfaces (Fig. 1D). For each backbone, we used ProteinMPNN and Rosetta FastRelax to design amino acid sequences optimized for interface complementarity and monomer stability, typically sampling eight sequence variants per backbone. We selected designs predicted with AlphaFold2 using initial-guess target templating (20) to fold as designed and bind FAK with high confidence (AF2 interface pAE scores < 10 and pLDDT > 90 for the complex, Fig. S1). After another round of partial diffusion of most promising designs, we selected 96 top-scoring designs for experimental testing. All designs were small single-domain, mostly helical proteins (∼100 amino acids).

Designed proteins were recombinantly expressed, purified using the semi-automated protein production (SAPP) pipeline (21) and their ability to express and monomeric behavior was tested on size-exclusion chromatography (SEC) (Fig. S2). We measured the binding affinity of 96 binders on SPR at a single concentration of 500 nM to select designs to move forward for closer characterization (Fig. S3). For SPR, the FAK kinase-domain target was site-specifically biotinylated with CoA-biotin and captured on a biotin-capture sensor chip, and purified designed proteins flowed over the immobilized kinase domain as analytes. We evaluated their activity in a high-throughput fluorescent kinase biosensor assay. In this assay, active FAK phosphorylates a substrate peptide between two fluorescent proteins that triggers a FRET decrease due to an SH2 domain that can only bind the phosphorylated peptide, while inhibited FAK maintains a high FRET signal (Fig. 1E). We incubated purified FAK with each designed protein and measured the kinase activity relative to a no-binder and small molecule inhibitor control thereby monitoring the time resolved FRET efficiency.

A number of the designs caused clear deviations in rate of FRET change (Fig. 1F). Three designs increase the rate of loss of fluorescence signal compared to FAK alone (suggesting they enhance FAK activity), whereas 30 other designs blocked FAK mediated FRET signal decrease (suggesting inhibition of FAK). The remaining designs had little to no effect or only modest perturbations of FAK activity. The minibinders FAK_A1 and FAK_A2 (green stars) increased the biosensor signal by approximately 150-200% compared to FAK alone, whereas FAK_I1 and FAK_I2 (red stars) reduced the signal to ∼20-30% of baseline, indicating activation and strong inhibition, respectively (Fig. 1F).

### Biochemical Characterization of FAK Activators and Inhibitors

We further characterized the activities of the four most potent binders using a series of biochemical assays. SPR was used to measure the binding affinity of each designed protein to the FAK kinase domain (Fig. 2AB). The four binders bound FAK with moderate to high affinity; the inhibitor designs FAK_I1 and FAK_I2 bound FAK with K_D_ values of 2.7 nM and 19 nM, respectively, while the activators FAK_A1 and FAK_A2 bound with K_D_ values of approximately 391 nM and 4.9 nM, respectively.

**Figure 2.**
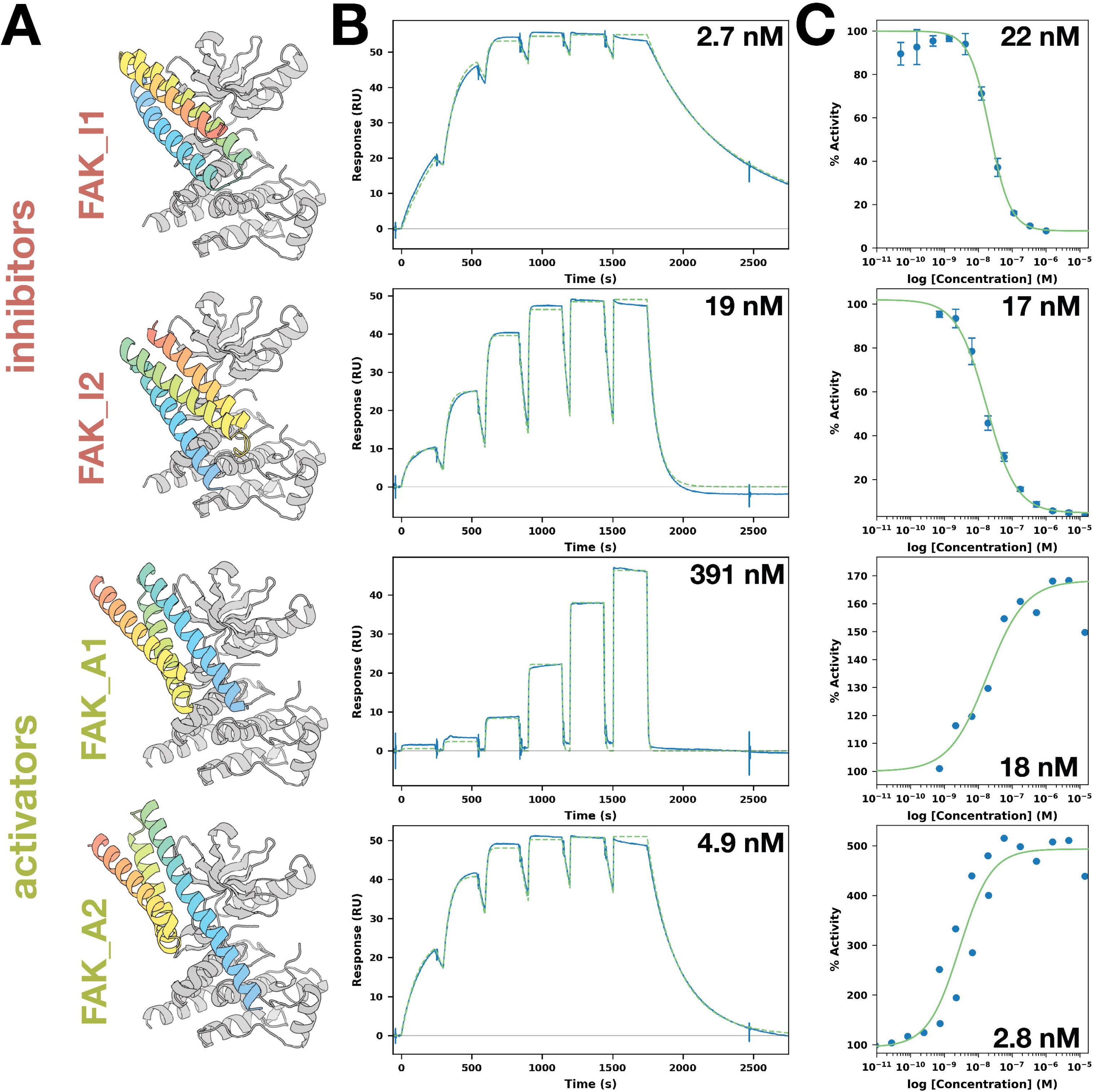
Biochemical characterization of *de novo* FAK modulators. **(A)** Models of the four top designs bound to the FAK kinase domain, highlighting the engineered interfaces of inhibitor binders FAK_I1 and FAK_I2 and activator binders FAK_A1 and FAK_A2. **(B)** Surface plasmon resonance sensorgrams showing binding of each designed protein to the FAK kinase domain. Solid lines represent experimental traces and dashed lines represent global fits. All four binders exhibit high-affinity, saturable interactions. **(C)** Radioactive kinase activity dose-response curves for inhibitor and activator designs. FAK_I1 and FAK_I2 potently inhibit FAK catalytic output with low-nanomolar IC50 values, whereas FAK_A1 and FAK_A2 increase FAK activity in a concentration-dependent manner. Together, these data confirm that the designed proteins function as strong inhibitors or activators of FAK in vitro.

We evaluated the effect of the designs on FAK catalytic activity using a radioactive kinase reaction assay (run by Reaction Biology) that quantifies incorporation of [γ-³²P] into a peptide substrate. The inhibitory designs produced a strong and consistent reduction in phosphate transfer. Concentration-response measurements yielded EC_50_ values of 22 ± 3 nM for FAK_I1 and 17 ± 3 nM for FAK_I2 (Fig. 2C), confirming potent inhibition in the low-nanomolar range, consistent with the affinities measured by SPR. In contrast, the activators increased radioactive phosphate incorporation. FAK_A1 (EC_50_ values of 18 ± 9 nM) approximately doubled the total ³²P incorporation compared to FAK alone, while FAK_A2 (EC50 values of 2.8 ± 0.9 nM) led to a more pronounced increase.

### Structural Basis of Allosteric Inhibition and Activation

To define the structural basis by which the designed binders modulate FAK activity, we solved a crystal structure of the FAK kinase domain bound to the activating minibinder FAK_A1 and compared it with the corresponding design model (Fig. 3AB). The co-crystal structure was refined to 2.04 Å resolution and closely recapitulated the designed binding mode. After superposition on the kinase domain, the binder backbone agreed with the model to ∼1.0 Å RMSD, while the binder fold itself overlaid with a backbone RMSD of 0.28 Å. Thus, the experimentally observed complex validates the intended binder geometry and shows that the designed protein can be accommodated on the kinase surface without a large-scale rearrangement of the kinase core.

**Figure 3.**
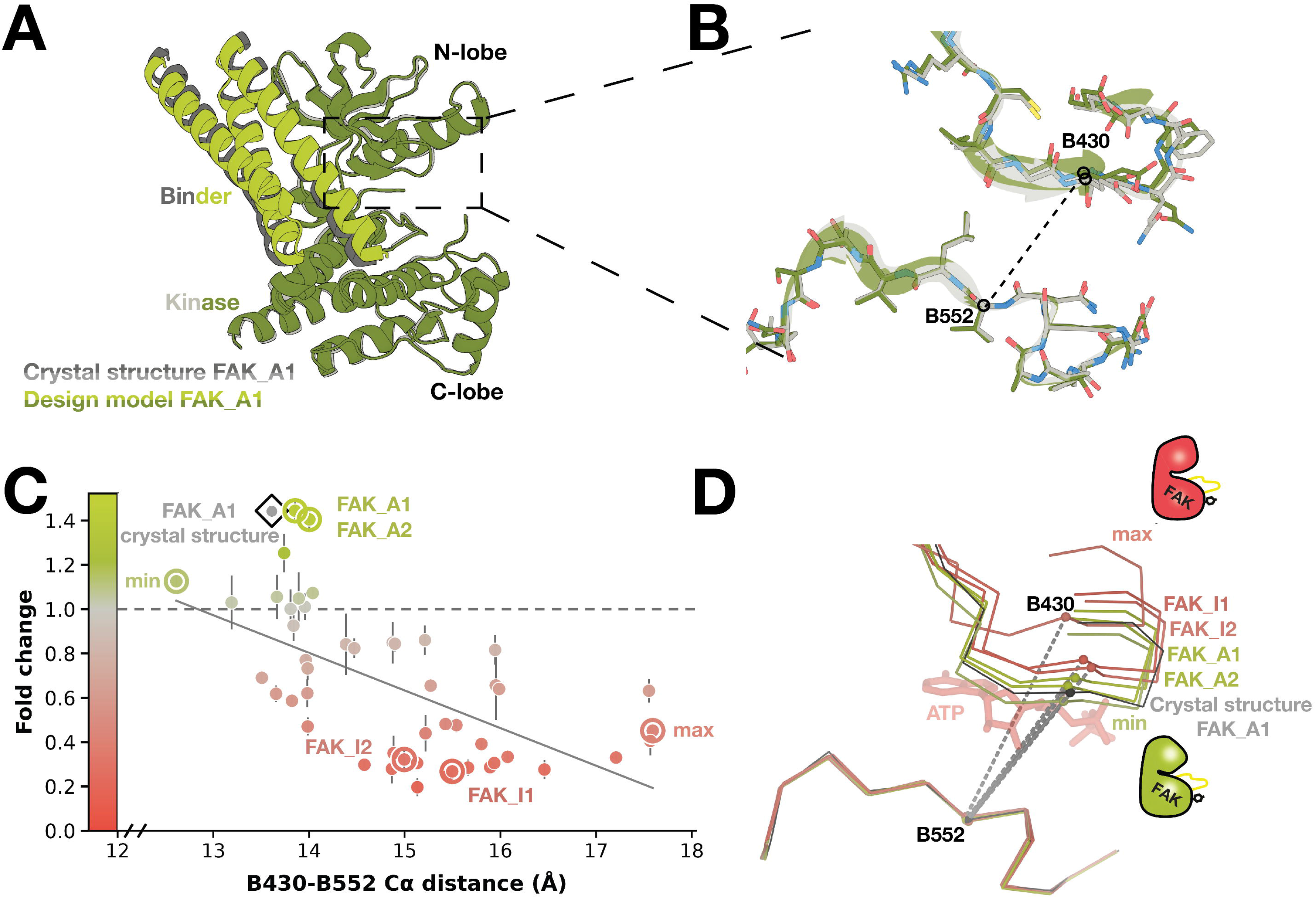
Structural analysis of inhibition and activation. **(A)** Overall view of the FAK_A1 complex, comparing the design model with the matched crystal structure after superposition on the kinase chain. The close agreement between the design model and crystal structure is evident across both the kinase and binder components. In the design model, the designed binder is shown in bright green and the kinase in dark olive. In the crystal structure, the kinase is shown in light grey and the binder in dark grey. **(B)** Magnified view of the same aligned complex, centered on the ATP-site-proximal region and shown in the same orientation as panel A. The kinase side-chains show close agreement between the design model and crystal structure, highlighting the accuracy of the modeled local side-chain geometry. Colors match panel A: dark olive for the design model and light grey for the crystal structure. The B430-B552 Cα-Cα distance used for the geometric analysis in panels C and D is indicated by a dashed grey connector. **(C)** Relationship between local ATP-pocket geometry and fold-change activity across the AF3-predicted ensemble. The x-axis reports the absolute B430-B552 Cα distance, with models aligned to their C-lobes. Points represent AF3 models, vertical bars show SEM of fold change, and color indicates activity fold change. The grey line shows a linear fit to the AF3 models (Pearson r = -0.60, n = 49). Highlighted designs are labeled as FAK_A1, FAK_A2, FAK_I1, and FAK_I2; the matched FAK_A1 crystal structure is shown as a white diamond with a black outline and center point. **(D)** Structural overlay of FAK_A1, FAK_A2, FAK_I1, FAK_I2, and the FAK_A1 crystal structure after the same C-lobe alignment. Cα traces for residues B425-B435 and B546-B558 are shown, with dashed grey segments connecting the measured B430 and B552 Cα positions. ATP from 2IJM is shown as reference in transparent red sticks to mark the nucleotide-binding site. Together, these data support a model in which inhibitory/activator designs are associated in altering the local ATP-pocket geometry.

FAK_A1 engages both the N- and C-lobe of the kinase domain as well as the hinge region connecting the two lobes. This binding mode positions the minibinder across structural elements that frame the ATP-binding site, suggesting that activity modulation may arise from altered local pocket geometry set by different placements of N- and C-lobe relative to each other rather than from global unfolding or overall distortion of the kinase. Consistent with this model, after alignment on the C-lobe, the FAK_A1 crystal structure and the AF3 models of FAK_A1, FAK_A2, FAK_I1, and FAK_I2 show close superposition of the C-lobe core (pairwise Cα RMSDs = 0.15-0.30 Å), whereas their N-lobes occupy different positions relative to the C-lobe, resulting in distinct ATP-binding-pocket geometries.

We used the experimentally validated FAK_A1 complex as an anchor for interpreting the remaining activating and inhibitory designs. To relate structural variation to functional output, we aligned AF3 models of the predicted complexes with the FAK_A1 crystal structure over the kinase C-lobe residues B507-B563 and B587-B686, excluding the more variable B564-B586 segment. We then quantified local ATP-pocket geometry using the distance between B430 Cα and B552 Cα relative to a 2J0L-derived reference. This simple geometric metric captured a functional trend across the design set: activating designs FAK_A1 and FAK_A2 remained close to the reference-like pocket geometry, whereas inhibitory designs FAK_I1 and FAK_I2 showed larger positive distance shifts. Across the designs, the B430-B552 distance was negatively correlated with fold change (Pearson r = -0.60), indicating that expansion or displacement of this local pocket measure is associated with reduced activity (Fig. 3CD).

Together, the crystal structure and ensemble analysis support a model in which designed binders tune FAK activity by stabilizing distinct local ATP-pocket conformations. In this model, FAK_A1 preserves a geometry compatible with activation, whereas inhibitory binders bias the ATP-site-proximal region toward a more open or displaced configuration. The structural agreement between the FAK_A1 crystal structure and the design model strengthens this interpretation by showing that the designed binding pose is experimentally realized and can be used to rationalize the activity-dependent conformational differences observed across the design series.

### Selectivity of Designed Kinase Modulators

An essential consideration for any kinase modulator is target selectivity. Kinases share conserved folds and often similar active-site chemistries, so off-target binding can lead to unintended pathway perturbation (3). Because FAK_I1 and FAK_I2 were designed to recognize nonconserved allosteric surfaces on FAK, we anticipated that they would preferentially inhibit FAK over other kinases. To assess this, both binders were tested at 2 µM against a representative set of kinases consisting of FAK, IRAK4, EGFR/ErbB1, CSNK1G1/CK1γ1, ROCK1, GSK3B/GSK3β, SYK, MKNK1/MNK1, MAPKAPK2 and ZAP70 (Fig. 4). Both binders strongly inhibited FAK, whereas inhibition of the other kinases was substantially lower to negligible, including for the tyrosine kinases EGFR, SYK and ZAP70 which are close relatives of FAK. The assay concentration was approximately 90-fold above the FAK_I1 IC₅₀ and 120-fold above the FAK_I2 IC₅₀; even at concentrations far exceeding those required for half-maximal FAK inhibition, both designed binders retained a pronounced preference for FAK over the other kinases tested.

**Figure 4.**
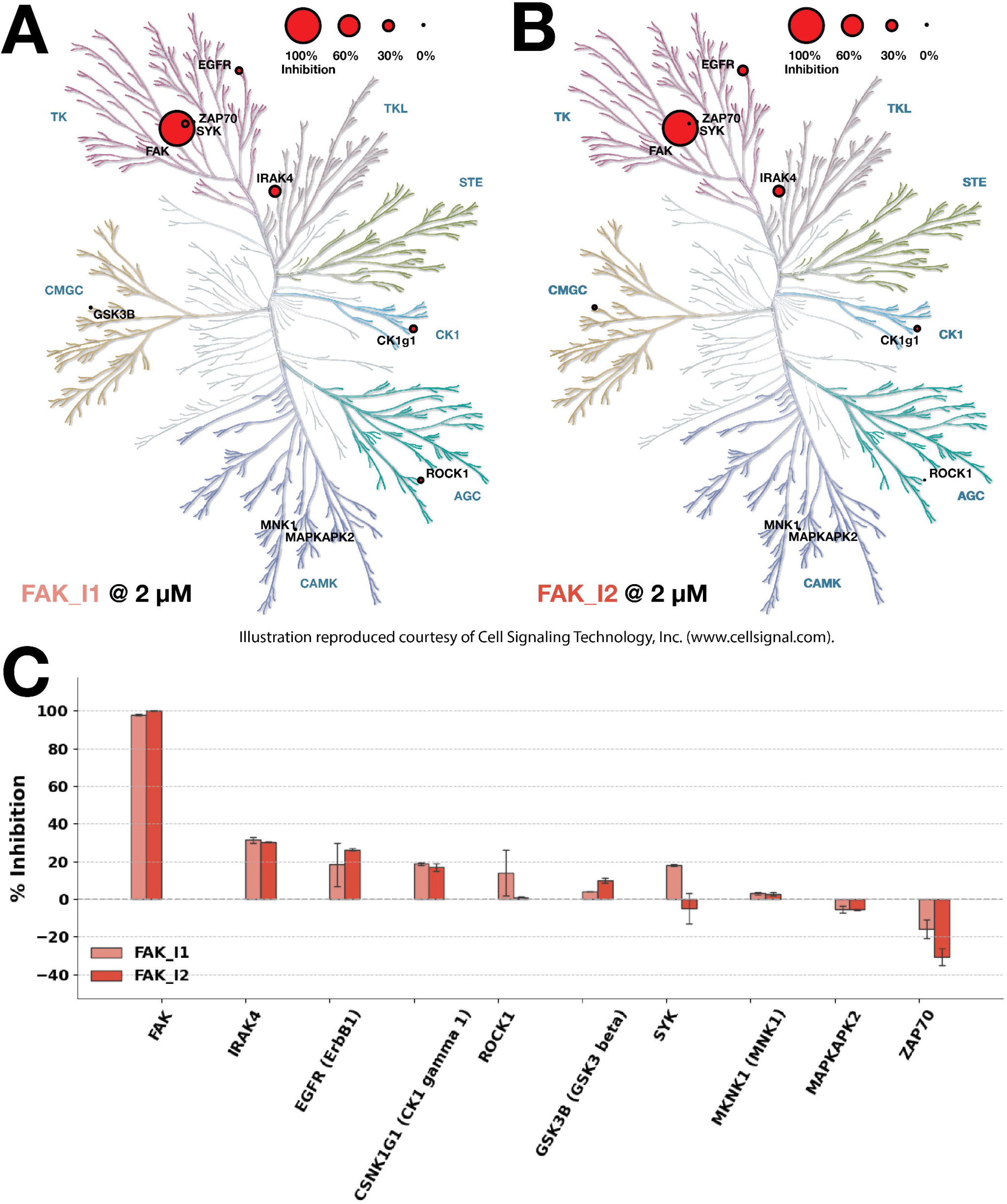
Kinase selectivity profile of the *de novo* designed FAK modulators. Mean percentage inhibition produced by FAK_I1 (**A**) and FAK_I2 (**B**), each tested at 2 µM, mapped onto the human kinome dendrogram. Circle size indicates the degree of inhibition; only kinases included in the biochemical panel are marked. Both designs strongly inhibit FAK while producing negligible inhibition of the other kinases tested. Illustration reproduced courtesy of Cell Signaling Technology, Inc. (www.cellsignal.com), with larger circles indicating stronger inhibition. FAK_I1 and FAK_I2 exhibit robust inhibition of FAK and minimal activity across the rest of the kinome, including closely related tyrosine kinases. **(C)** Mean % inhibition of FAK_I1 (salmon) and FAK_I2 (dark red) across the kinase panel, with error bars showing half-range between replicate values and targets ordered by overall inhibition.

### Designed kinase binders engage and modulate kinase signaling in cells

FAK is commonly overexpressed, amplified, and hyperactivated in several cancers including triple-negative breast cancer (TNBC), ovarian cancer and pancreatic ductal adenocarcinoma (PDAC) (22). The FAK inhibitor defactinib, in combination with the FAK-MEK clamp inhibitor (avutometinib) has been recently FDA-approved for the treatment of *KRAS*-mutated low-grade serous ovarian cancer, highlighting the therapeutic potential of disabling FAK in cancer (22). Our prior work has demonstrated that FAK inhibition or targeted degradation disrupts biological responses in TNBC and PDAC cells (23, 24). To study the *de novo* FAK modulators in cells and determine whether we could control FAK activity and cellular outcomes, we first evaluated expression in MDA-MB-231 cells, a TNBC cell line with amplified FAK and dependence on FAK in 3D-spheroid models (23, 24). To stably express the *de novo* FAK modulators in cells, we designed lentiviral constructs to transduce MDA-MB-231 cells and enriched for highly expressing pools (Fig. S5 AB). Following successful integration and expression, we confirmed cellular engagement of FAK by evaluating whether the *de novo* modulators could prevent PROTAC-mediated FAK degradation (23). BSJ-04-146 is a FAK PROTAC composed of a selective FAK inhibitor that binds the FAK active site conjugated to a cereblon binding ligand (23). We observed that the expression of each *de novo* FAK modulator prevented FAK degradation by BSJ-04-146, indicating FAK engagement in cells (Fig. 5A). Levels of rescue were equivalent to controls in which MDA-MB-231 cells were pretreated with the NEDD8 inhibitor MLN4924 or the FAK inhibitor BSJ-04-175. Next, we evaluated whether the *de novo* FAK modulators altered FAK activity by monitoring the phosphorylation status of FAK at Y397, its primary autophosphorylation site (25, 26). As expected, BSJ-04-146 treatment reduced both total and phosphorylated FAK, whereas treatment with the FAK inhibitor defactinib reduced phosphorylated FAK without affecting total FAK levels (Fig. 5B). Similar to our in vitro findings (Fig. 2), MDA-MB-231 cells expressing FAK_I1 and FAK_I2 reduced phosphorylated FAK, while cells expressing FAK_A1 and FAK_A2 increased phosphorylated FAK (Fig. 5B). All four *de novo* FAK modulators modestly increased levels of FAK protein, suggesting that binding, whether inhibitory or activating, may also increase the stability of FAK.

**Figure 5.**
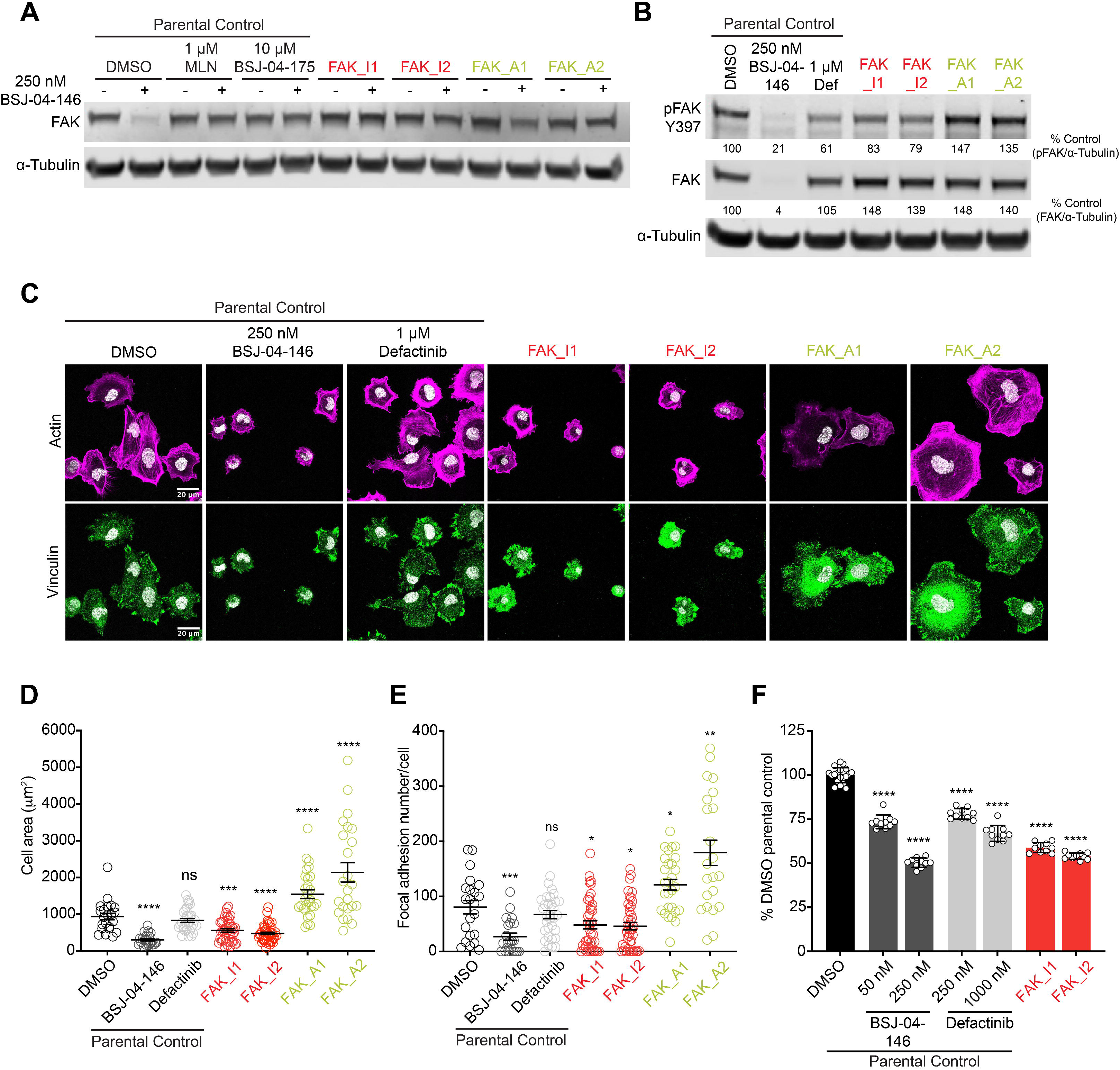
Cellular characterization of *de novo* FAK modulators. **(A)** Immunoblot analysis of the indicated MDA-MB-231 cells. MDA-MB-231 parental control cells were pretreated with DMSO, 1 μM MLN4924 (MLN), or 10 μM BSJ-04-175 for 2 hours prior to the treatment of 250 nM BSJ-04-146 for an additional 4 hours. MDA-MB-231 cells stably expressing FAK_I1, FAK_I2, FAK_A1 or FAK_A2 were treated with DMSO or 250 nM BSJ-04-146 for 4 hours. **(B)** Immunoblot analysis of the indicated MDA-MB-231 cells. MDA-MB-231 parental control cells were treated with DMSO, 250 nM BSJ-04-146 or 1 μM defactinib for 24 hours. MDA-MB-231 cells stably expressing FAK_I1, FAK_I2, FAK_A1 or FAK_A2 were treated with DMSO for 24 hours. **(C)** Representative images of cell spreading after MDA-MB-231 cells were plated for 60 min on a collagen coated surface. MDA-MB-231 parental control cells were treated with DMSO, 250 nM BSJ-04-146, or 1 μM defactinib for 24 hours prior to assessment for cell spreading. MDA-MB-231 cells stably expressing FAK_I1, FAK_I2, FAK_A1 or FAK_A2 were treated with DMSO for 24 hours prior to assessment for cell spreading. **(D-E)** Mean cell area **(D)** and mean number of focal adhesions per cell **(E)** were calculated from >15 cells in each condition shown in C. Data in A-E are representative of *n* = 3 independent experiments. ****P < 0.0001, ***P < 0.001, **P < 0.01, *P < 0.05, or not significant (ns) are indicated after analysis using a two-tailed Mann-Whitney test. **(F)** DMSO-normalized antiproliferation of the indicated MDA-MB-231 cells cultured as ultra-low adherent 3D spheroid suspensions for 120 hours. MDA-MB-231 parental control cells were treated with DMSO or the indicated doses of BSJ-04-146 or defactinib. MDA-MB-231 cells stably expressing FAK_I1 or FAK_I2 were treated with DMSO. Data in F are presented as the mean ± SD of 10-20 biologically independent samples and are representative of *n* = 3 independent experiments. ****P < 0.0001 is indicated after analysis using 1-way ANOVA with post hoc Dunnett’s test.

The changes in phosphorylation status of FAK prompted our evaluations of the biological consequences of the *de novo* modulators on FAK-dependent phenotypes. FAK is a critical kinase downstream of integrin signaling that controls cell spreading, adhesion, and migration and viability in 3D-models (22, 27). First, we evaluated whether each *de novo* FAK modulator could impede or enhance the spreading of MDA-MB-231 cells, as well as the formation of focal adhesions, upon plating onto collagen-I coated surfaces. Collagen-I is a major component of the extracellular matrix, and cell spreading on a collagen-I-coated surface models the initial attachment of cancer cells to the stromal microenvironment which requires FAK activity. This biological process is important in early events that promote metastatic dissemination, including cell adhesion, spreading, and acquisition of migratory behavior (28, 29). To test this phenotype, we imaged single cells using confocal microscopy and evaluated actin staining to quantify cell area and monitor cell spreading and vinculin staining to quantify focal adhesions (26, 30). Treatment with BSJ-04-146 or expression of FAK_I1 or FAK_I2 in MDA-MB-231 cells markedly reduced cell spreading and decreased the number of focal adhesions per cell (Fig. 5CE). At the evaluated dose of defactinib (1000 nM), we did not observe a statistically significant reduction in the cell spreading or focal adhesion numbers. Acute chemical inhibition versus prolonged inhibition by the constitutively expressed inhibitory *de novo* FAK modulators may contribute to these observed differences. By contrast, MDA-MB-231 cells expressing FAK_A1 or FAK_A2 displayed enhanced cell spreading and increased focal adhesion numbers (Fig. 5CE). This indicates that the *de novo* FAK modulators can toggle FAK activity to disrupt or enhance its functions in TNBC cells.

Next, we focused on the inhibitory *de novo* modulators, by examining MDA-MB-231 proliferation as 3D-spheroids, which is an assay that is strongly predictive of in vivo efficacy (23, 31). Consistent with our prior work, BSJ-04-146 or defactinib treatment led to an antiproliferative effect, with more potent and pronounced reductions with BSJ-04-146 (Fig. 5F). Expression of FAK_I1 and FAK_I2 in MDA-MB-231 cells significantly reduced the cell viability of 3D-spheroids (Fig. 5F). To extend these findings in an independent model, we expressed the inhibitory *de novo* FAK modulators in PATU-8988T cells (Fig. S6A), a PDAC cell line that we have previously shown is sensitive to FAK disruption when cultured as 3D-spheroids (23). FAK_I1 or FAK_I2 expression led to substantial reductions in the levels of phosphorylated FAK and significantly diminished viability of 3D-spheroids, to levels similar to BSJ-04-146 and defactinib (Fig. S6BC). Together, these findings indicate that the *de novo* FAK modulators effectively engage FAK in cells, alter its activity, and modify downstream cellular outcomes.

Having established that the designed binders modulate endogenous FAK signaling in cells, we asked whether the same design strategy could be extended to another kinase. We focused on Src kinase, a proto-oncogenic tyrosine kinase that, like FAK, contributes to integrin signaling, cancer cell motility and invasive behavior, but is regulated through a distinct architecture involving SH2- and SH3-domain interactions and inhibitory C-terminal phosphorylation. We reasoned that designed binders targeting regulatory surfaces on Src could similarly stabilize functional kinase states and thereby inhibit or enhance Src-dependent signaling output.

To generate Src-directed modulators, we used inactive Src structures, including SH2-bound conformations, as design targets. Starting from successful FAK inhibitory binders, we applied RFdiffusion with partial diffusion settings with Src substituted as the target, allowing the FAK inhibitor scaffolds to serve as structural templates for rapid retargeting to Src (32). The partially diffused backbones were then subjected to ProteinMPNN for sequence design against the Src target, and designs predicted by AlphaFold2 to form favorable Src-binding interfaces were selected for experimental testing (Fig. S4).

We tested the Src modulators in HEK293 cells using an intracellular phosphorylation assay. Cells were co-transfected with a constitutively active Src fusion, containing either full-length Src (FL) or a truncated kinase-domain construct (TRC), and a CD247 ITAM reporter. Each Src construct was tethered to either a designed binder or a nonphosphorylatable pseudosubstrate control. Phosphorylation was measured by flow cytometry using the pan-phosphotyrosine antibody pY100 and an antibody specific for CD247 pY142. The pY100 (XXX**Y**XXX) signal provides a broad, non-exhaustive measure of cellular phosphotyrosine abundance, whereas CD247 pY142 (HDGL**Y**QGLSTATKDTYDALHMQ) reports phosphorylation of a defined site on the ectopic reporter. Lower values in either assay therefore indicate stronger suppression of Src-associated phosphorylation. For each readout and Src context, the corresponding PPP2CA pseudosubstrate control construct (HITQVFGFFDE) was used as a benchmark.

Suppression varied across designs and Src contexts. Src_I1 produced the strongest and most consistent effect (Fig. S7). In the TRC context, both pY100 and CD247 pY142 signals were comparable to or lower than the corresponding PPP2CA benchmarks, and in the FL context Src_I1 produced the lowest signals among the designed fusions. Src_I2 also reduced both phosphorylation readouts in both Src contexts. Src_I3 and Src_I4 showed substantially stronger suppression with TRC than with FL Src, indicating context-dependent activity. Src_I6 produced an intermediate phenotype, whereas Src_I5 and Src_I7 retained relatively high phosphorylation signals and showed little suppression.

Together, these results establish that the designed binders function as intracellular kinase modulators rather than passive binding reagents. In cells, the FAK binders engaged endogenous FAK, protected it from PROTAC-mediated degradation, and altered Tyr397 phosphorylation and downstream adhesion phenotypes in a manner consistent with their inhibitory or activating activities. Retargeting the inhibitory scaffolds to Src similarly produced binders that suppressed both global tyrosine phosphorylation and phosphorylation of the defined Src substrate CD247. Thus, the design framework is modular and transferable, enabling genetically encoded binders to engage distinct kinase regulatory surfaces and tune signaling output in cells.

## Discussion

We show that *de novo* designed proteins can either inhibit or enhance kinase activity by engaging conformationally sensitive surfaces outside the conserved ATP-binding site. Using RFdiffusion we generated compact binders that either inhibited or enhanced FAK catalytic activity based on the input structure chosen, with the strongest inhibitors acting at low-nanomolar concentrations. The close agreement between the FAK_A1 design model and the 2.04-Å crystal structure validates the intended binding geometry and demonstrates that diffusion-based design can generate kinase interfaces with near-atomic accuracy. Structural comparisons further suggest that the binders modulate activity by altering ATP-site-proximal geometry rather than by stabilizing a single canonical DFG/αC-helix state or recreating the multidomain autoinhibited conformation of FAK. Activating binders preserved a more compact, active-like pocket geometry, whereas inhibitory binders were associated with a more open or displaced configuration. Because productive catalysis requires coordinated positioning of the glycine-rich loop, activation segment, αC helix and catalytic core, controlling the relative arrangement of these elements may provide a general route for engineering kinase activity.

The designed proteins function in cells as kinase activity modulators. Expression of the FAK binders protected endogenous FAK from PROTAC-mediated depletion to varying degrees, providing an orthogonal measure of target engagement in cells. The inhibitory binders reduced FAK Tyr397 phosphorylation and produced cellular phenotypes associated with impaired FAK signaling, including reduced spreading, fewer vinculin-positive focal adhesions and decreased viability signals. Conversely, the activating binders increased cell spreading and focal-adhesion abundance. They did not further increase the basal pY397-FAK/FAK ratio, potentially because the accessible pool of FAK that can be phosphorylated at Tyr397 was already near saturation under these conditions. The opposing adhesion phenotypes therefore provide a functional readout of bidirectional modulation that is not captured by Tyr397 phosphorylation alone.

Retargeting the inhibitory scaffolds from FAK to Src further demonstrates the adaptability of the approach. Partial diffusion allowed the overall scaffold architecture to be retained while remodeling the interface for a distinct kinase surface, and the resulting Src-directed proteins suppressed both global tyrosine phosphorylation and phosphorylation of the defined Src-family substrate CD247 in cells. Although broader testing across kinases will be required to determine how generally scaffolds can be transferred, these results show that successful modulator backbones can serve as starting points for rapid interface redesign rather than requiring an entirely new design campaign for every target.

Synthetic kinase activators have been particularly difficult to generate because activation requires increasing the occupancy of catalytically productive states rather than simply blocking substrate or ATP binding. Previous work with engineered monobodies established that protein scaffolds can allosterically alter kinase conformational equilibria. The present results extend this principle to *de novo* proteins and show that inhibitory and activating functions can emerge from the same structure-guided workflow. Genetically encoded modulators could therefore complement small-molecule inhibitors by enabling sustained, cell-type-specific or spatially restricted perturbation of kinase activity. Their compact size also makes them suitable for fusion to localization domains, inducible systems or degradation machinery.

The ability to design selective, genetically encoded inhibitors and activators against defined kinase surfaces provides a framework for dissecting signaling mechanisms and engineering cellular behavior. For therapeutic applications, delivery, immunogenicity and comprehensive specificity will be important considerations. More generally, the results suggest that conformationally sensitive regulatory surfaces can be treated as programmable design targets for controlling enzyme activity in cells.

## Materials and Methods

### Protein binder design for FAK activation and inhibition

FAK kinase-domain structures (PDB IDs 2J0L, 4D4R, 4D5H, 4K9Y, 1MP8 and 4D55) were used as conformational target templates. Solvent-exposed surfaces were selected as design sites, with targeted hotspot residues listed in Supplementary Table T1.

Binder backbones of 80-120 residues were generated against fixed FAK structures using the RFdiffusion protein-protein interface design protocol (33). For each backbone, eight sequences were generated with ProteinMPNN and refined using Rosetta FastRelax. Designs were evaluated using the AlphaFold2 initial-guess protocol described by Bennett et al. (20) and selected based on recovery of the intended binder fold and interface, high pLDDT and an interface predicted aligned error below 5 Å. Top candidates underwent an additional round of RFdiffusion partial diffusion (32), sequence design and structural filtering. Ninety-six designs were selected for experimental characterization.

For structure-activity analysis, selected binder-FAK complexes were repredicted without constraining the FAK coordinates, using Alphafold3 (34) using a multiple-sequence alignment for FAK and a single sequence for the designed binder. The resulting structural ensembles were used to assess binder-associated changes in kinase conformation.

### Minibinder expression and purification by Semi-Automated Protein Production (SAPP)

Minibinder coding sequences were synthesized as linear, double-stranded eBlocks™ Gene Fragments (Integrated DNA Technologies, IDT) containing terminal BsaI recognition sites and compatible Golden Gate overhangs. The fragments were supplied in 384-well PCR plates at 10 ng µL⁻¹ in 20 µL per well, in nuclease-free water or IDTE buffer, pH 8.0, and were used directly for cloning without prior PCR amplification.

Minibinder sequences were cloned into the pET29b-derived LM627 vector, which encodes a C-terminal SNAC-His₆ affinity tag and contains a ccdB negative-selection cassette between the BsaI cloning sites. Golden Gate assembly reactions were prepared in 96-well PCR plates using an Echo 525 acoustic liquid handler (Beckman Coulter). Each 1-µL reaction contained 0.1 µL 10× T4 DNA ligase buffer, 1.2 U BsaI-HFv2, 40 U T4 DNA ligase, 2-4 fmol LM627 vector, and a twofold molar excess of the corresponding eBlock insert. Reactions were incubated at 37 °C for 20 min, followed by 60 °C for 5 min. Assembly products were transformed directly without intermediate colony isolation or plasmid preparation.

For transformation, 1 µL of each Golden Gate reaction was combined with 6 µL chemically competent Escherichia coli BL21(DE3) cells in a 96-well PCR plate. Cells were incubated on ice for 30 min, heat-shocked at 42 °C for 10 s, returned to ice for 2 min, and recovered in 100 µL SOC medium for 1 h at 37 °C with orbital shaking at 1,000 rpm. Each recovered transformation was divided among four 1-mL expression cultures in round-bottom 96-deep-well plates, giving a total expression volume of 4 mL per minibinder. Expression medium consisted of TBII supplemented with 5 g L⁻¹ glycerol, 0.5 g L⁻¹ glucose, 2 g L⁻¹ lactose, 2 mM MgSO₄, and 50 µg mL⁻¹ kanamycin. Plates were covered with gas-permeable seals and incubated for at least 20 h at 37 °C with shaking at 1,000 rpm. Cells were harvested by centrifugation at 4,000 × g for 5 min, and the culture supernatants were discarded.

Cell pellets were resuspended in 100 µL lysis buffer per millilitre of expression culture. The lysis buffer consisted of B-PER bacterial protein extraction reagent supplemented with 0.1 mg mL⁻¹ lysozyme, 25 U mL⁻¹ Benzonase, and 1 mM phenylmethylsulfonyl fluoride. Lysis was performed for 15 min at 37 °C with agitation at 1,000 rpm. The four lysates corresponding to each minibinder were consolidated into a single well, and insoluble material was removed by centrifugation at 4,000 × g for 15 min.

His-tagged minibinders were purified from the clarified soluble fraction by immobilized-metal affinity chromatography in a 96-well plate format. Ni-NTA resin was distributed into 96-well fritted plates at a bed volume of 50 µL per well and equilibrated with wash buffer containing 20 mM Tris-HCl, pH 8.0, 300 mM NaCl, and 25 mM imidazole. Clarified lysates were applied to the resin and allowed to pass through under low vacuum. The resin was washed three times with 400 µL wash buffer. Bound minibinders were eluted with 200 µL of 20 mM Tris-HCl, pH 8.0, 300 mM NaCl, and 500 mM imidazole. Eluates were sterile-filtered by centrifugation through a 0.22-µm 96-well filter plate.

Filtered IMAC eluates were analyzed and further purified by size-exclusion chromatography using a Superdex 75 5/150 GL column operated at up to 0.65 mL min⁻¹ in 20 mM sodium phosphate, pH 7.4, and 100 mM NaCl. Protein elution was monitored by absorbance at 280 nm, and fractions corresponding to the principal monomeric peak were collected. Protein yields were calculated from the integrated 280-nm absorbance using sequence-derived molecular masses and extinction coefficients. Apparent molecular masses and oligomeric states were estimated by comparison of SEC retention volumes with a column calibration curve. Fractions containing soluble, predominantly monomeric minibinders were pooled for subsequent SPR measurements.

### FAK biotinylation and surface plasmon resonance

The purified FAK kinase-domain construct, containing an N-terminal His₆-ybbR tag and a C-terminal Strep-tag II, was site-specifically biotinylated using Sfp phosphopantetheinyl transferase and CoA-biotin. FAK was supplied at 2.02 mg mL⁻¹ (55.3 µM) in 20 mM HEPES, pH 7.0, 150 mM NaCl, 5% glycerol, and 2 mM TCEP. For each labeling reaction, 25 µL FAK (1.38 nmol) was combined with 1.38 µL of 5 mM CoA-biotin (6.92 nmol; fivefold molar excess), 3 µL Sfp to approximately 5 µM final concentration, and 3 µL of 10 mM MgCl₂ to approximately 1 mM final concentration, yielding a total reaction volume of 32.4 µL. The reaction was incubated for approximately 5 h at room temperature. The sample was subsequently diluted with 20 µL of 1× HBS-EB running buffer, and unreacted CoA-biotin and other low-molecular-weight components were removed using a Zeba spin-desalting column equilibrated in HBS-EB. The concentration of the recovered biotinylated FAK was determined by absorbance at 280 nm using a calculated molecular mass of 36,492 Da and an extinction coefficient of 47,900 M⁻¹ cm⁻¹. The recovered protein concentration was 0.7505 mg mL⁻¹, corresponding to 20.57 µM.

Surface plasmon resonance measurements were performed on a Biacore 8K by Cytiva using a biotin-capture sensor chip (28920233). HBS-EB running buffer was prepared by diluting a 10× stock tenfold with Milli-Q water and was filtered and degassed before use. Biotinylated FAK was diluted in running buffer to 3.75 µg mL⁻¹, corresponding to approximately 103 nM, and captured on the active sensor surface. A reference surface without captured FAK was used to correct for nonspecific surface binding and bulk refractive-index changes.

For the primary binding screen, purified designed proteins were diluted to 500 nM in HBS-EB, with 130 µL prepared per sample, and injected as soluble analytes over the immobilized FAK kinase domain. Lead binders were subsequently characterized using a twofold concentration series with a maximum analyte concentration of 2 µM. Kinetic measurements were performed in a single-cycle format, with sequential analyte injections without surface regeneration between concentration steps, followed by a final dissociation phase. Sensorgrams were double-referenced by subtracting both the reference-channel response and the response from buffer-only injections. Corrected sensorgrams were globally fit to determine the association rate constant (kₐ), dissociation rate constant (kd), and equilibrium dissociation constant (KD = kd/kₐ).

### Biosensor screening assay

FAK activity was measured using an intramolecular ECFP-SH2-YPet FRET biosensor adapted from Seong et al. (35). Each 19 µL reaction contained 260 nM FAK, 700 nM FRET biosensor, 1.3 µM of the indicated minibinder, and 120 µM ATP. A reaction containing 5 µM staurosporine served as the negative kinase-activity control. FAK-containing reactions were incubated with the minibinders for 1 h before initiation of the kinase reaction by ATP addition. A reaction containing 5 µM staurosporine instead of minibinder was treated identically and served as the negative kinase-activity control. A reaction containing FAK, biosensor, ATP, and vehicle without minibinder or staurosporine served as the positive activity control, while a reaction lacking ATP served as an assay-background control. ATP was added sequentially to the wells row-wise at 30-s intervals to be able to accurately measure activity. Fluorescence was measured kinetically at 2-min intervals using a Synergy Neo2 multimode microplate reader. Samples were excited at 430-433 nm, and ECFP and YPet emissions were measured at 526-527 nm, respectively. FAK activity was quantified from the time-dependent increase in the ECFP/YPet emission ratio. Reactions were always performed in triplicate and repeated twice with a repeated expression of minibinders.

### Radioactive kinase activity assay

Kinase activity was measured by Reaction Biology Corp. (Malvern, PA, USA) using the HotSpot radiometric kinase assay (36). Test minibinders FAK_I1, FAK_I2, FAK_A1 and FAK_A2 were assayed against FAK in 10-point concentration-response format using 3-fold serial dilutions. Reactions were carried out at 1 µM ATP, and staurosporine was included as a control inhibitor in a 10-point, 4-fold dilution series starting at 20 µM. Kinase activity was quantified from radiolabeled phosphate incorporation into substrate and normalized to DMSO-treated controls to yield percent activity. Counter-assay reactions, in which test articles were added immediately before transfer to P81 paper, were run in parallel to assess assay interference.

Concentration-response data were replotted from the Reaction Biology reports and fit with four-parameter logistic models. FAK_I1 and FAK_I2 were analyzed as inhibitors and reported as IC50 values, whereas FAK_A1 and FAK_A2 increased FAK activity and were analyzed as activators with EC50 values. Error bars represent standard deviation for triplicate measurements.

### Kinase selectivity profiling

Kinase selectivity profiling was performed by Thermo Fisher Scientific using the SelectScreen Z′-LYTE biochemical kinase assay service. FAK_I1 and FAK_I2 were tested at 2 µM against FAK (PTK2), IRAK4, EGFR/ErbB1, CSNK1G1/CK1γ1, ROCK1, GSK3B/GSK3β, SYK, MKNK1/MNK1, MAPKAPK2 and ZAP70 under kinase-specific assay conditions. The Z′-LYTE assay uses a doubly labelled FRET peptide substrate. Following the kinase reaction, a development protease selectively cleaves non-phosphorylated peptide, disrupting FRET, whereas phosphorylated peptide remains intact. Kinase activity was quantified from the ratio of coumarin emission at 445 nm to fluorescein emission at 520 nm after excitation at 400 nm. Percentage inhibition was calculated relative to active-kinase and no-ATP controls according to the provider’s standard protocol.

### Co-Crystalization and structure determination

The avian FAK kinase domain (residues 411-686) was expressed and purified as described previously (15, 16). The FAK:FAK_A1 complex was setup at 7 mg/ml FAK kinase with a 9 fold molar excess of FAK_A1. Various sparse matrix crystallisation screens were setup in a sitting drop format by mixing equal volumes of protein complex and precipitant solution. Best crystals appeared in condition B2 of Jene Biosciences classic 2 screen, containing 16% PEG 4000, 200 mM Sodium acetate and 100 mM Tris pH8.5. Crystals were frozen in liquid nitrogen in crystallisation condition with 20% Glycerol as cryoprotectant. Diffraction data was collected at the XALOC-BL13 beamline of the ALBA synchrotron (Barcelona, Spain) at 100 K using a PILATUS 6 M detector. Diffraction data was indexed and integrated with XDS (37) and scaled with AIMLESS (38), applying a high resolution cut-off of 2.04 Å. Crystals belong to the P22121 spacegroup with unit cell parameters a=42.22 Å, b=87,86 Å, c=109,94 Å and all angles 90°. Phases were determined by molecular replacement using PHASER (39), using the structure of the FAK kinase domain (PDB: 1MP8) as search probe. Refinement and model building was performed by iterative cycles of restrained refinement, including TLS cycles, with REFMAC5 (40) and manual building guided by electron density maps, using COOT (41, 42). Final R-factors are 23.8/26.6 (work/free) (for full crystallographic table, see supplementary Table S1).

### Cell line culturing

MDA-MB-231 (ATCC, #HTB-26), PATU-8988T (DSMZ, #ACC-162), and 293FT (Thermo Fisher Scientific #R70007) cells were cultured in Dulbecco’s modified Eagle’s medium (DMEM) supplemented with 10% fetal bovine serum (FBS), 100 U/mL penicillin, and 100 μg/mL streptomycin at 37 °C with 5% CO_2_. Cell lines were routinely confirmed to be mycoplasma negative using the MycoProbe Mycoplasma detection kit (R&D systems, #CUL001B).

### Plasmid generation

The nucleotide sequences encoding FAK modulators (FAK_I1, FAK_I2, FAK_A1, or FAK_A2) were codon-optimized, cloned in-frame with T2A-EGFP into the pGenLenti vectors, and synthesized by GenScript (Piscataway, NJ, USA). Plasmids were amplified in NEB Stable E. coli cells (NEB, #C3040H) with selection on LB agar plates containing carbenicillin (100 μg/mL). Individual colonies were isolated, and all constructs were sequence-verified.

### Viral production and stable cell line generation

Lentivirus was produced and concentrated as previously described (43). HEK293FT cells were co-transfected with pMD2.G (Addgene, #12259), psPAX2 (Addgene, #12260), and the lentiviral FAK modulator plasmid of interest, using Lipofectamine 2000 (Thermo Fisher Scientific, #11668019), according to the manufacturer’s instructions. Viral particles were collected 60 hours after transfection, filtered through a 0.45 μM membrane, and concentrated using 1:3 Lenti-X Concentrator (Takara Bio, #631231), according to the manufacturer’s instructions. MDA-MB-231 and PATU-8988T cells were transduced with concentrated lentiviral supernatant for 24 h in the presence of 4 μg/mL polybrene (Millipore Sigma, #TR1003G), followed by selection with 2.0 μg/mL puromycin.

### Fluorescence-activated cell sorting (FACS)

10 million MDA-MB-231 or PATU-8988T cells were trypsinized and pelleted by centrifugation at 1000 rpm for 5 min. The pellet was resuspended in 2.5 mL sorting buffer consisting of 1X PBS, 2 % FBS, and 0.5 mM EDTA. Following resuspension, samples were filtered through a polystyrene round bottom tube with cell strainer (Falcon, #28-154) and sorted for GFP expression using an MA900 Cell Sorter (Sony). For each cell line, 300,000 sorted cells were collected into polystyrene tubes with 2 mL culture medium, pelleted by centrifugation and resuspended in fresh medium, and transferred to 6-well tissue culture plates. To confirm GFP expression after sorting, phase contrast and GFP images were acquired using an ECLIPSE Ts2 inverted microscope (Nikon) equipped with a Digital Sight 10 camera (Nikon) and a 10X phase contrast objective.

### Compounds

Defactinib (MedChemExpress, #HY-12289) and MLN4924 (Calbiochem, #5054770001) were purchased from the indicated sources. BSJ-04-146 and BSJ-04-175 were synthesized as previously described (23).

### Immunoblotting

Immunoblotting was performed as previously described (31). Cells were lysed in RIPA buffer containing cOmplete protease inhibitor (Millipore Sigma, #11873580001), PhosSTOP phosphatase inhibitor (Millipore Sigma, #4906837001), and 0.04% Benzonase (Millipore Sigma, #712063) at 4 °C with gentle end-to-end rotation for 1 hour. Lysates were centrifuged at 15000 rpm for 10 minutes at 4 °C and clarified supernatants were transferred to pre-chilled tubes. Protein lysates were quantified using a Pierce BCA Protein Assay kit (Thermo Fisher Scientific, #23225) and loaded in equal amounts on 4–12% SDS-PAGE gradient gels, followed by transfer onto nitrocellulose membranes using an iBlot3 (Thermo Fisher Scientific) and overnight incubation with the primary antibody as indicated. The next day, membranes were washed with TBS-T (TBS with 0.1% Tween) and incubated with appropriate fluorescently labeled infrared secondary antibodies (LICOR, IRDye) at room temperature for 1 hour. Membranes were imaged using the LI-COR Odyssey Imaging System. The following primary antibodies were used in this study: FAK rabbit (Cell Signaling Technology, #13009), FAK mouse (Cell Signaling Technology, #62220), FAK phospho-Y397 (Cell Signaling Technology, #8556), and α-Tubulin (Cell Signaling Technology, #3873). Quantifications were performed using ImageJ as indicated.

### Analysis of cell viability in 3D-spheroid culture

3D-spheroid culture experiments were performed as previously described (23). MDA-MB-231 and PATU-8988T cells were plated in PrimeSurface 384-well 3D culture spheroid plates (S-bio, #MS-9384WZ). MDA-MB-231 cells were plated at a density of 500 cells per well in 20 μL media containing 10% Matrigel (Corning, #356231), and PATU-8988T cells were plated at a density of 200 cells per well in 50 μL media. The next day, cells were treated with compounds using a D300e digital dispenser (HP), and all wells were normalized to 200 nL total DMSO. Plates were incubated for 120 hours in a 5% CO_2_ incubator at 37 °C. For cell viability assessment, plates were equilibrated to room temperature. For MDA-MB-231 spheroids, 20 μL 3D CellTiter-Glo (Promega, #G9682) was added to each well. For PATU-8988T spheroids, 10 μL CellTiter-Glo (Promega, #G7570) was added to each well. After shaking at room temperature for 5 minutes and incubation at room temperature for 25 minutes, luminescence was measured on the CLARIOstar Plus Plate Reader (BMG LabTech). The data was normalized to DMSO-treated control wells and analyzed using GraphPad PRISM v10.

### Cell spreading on collagen and confocal microscopy

MDA-MB-231 cells were plated at a density of 100,000 cells in 2 mL in 6-well tissue culture plates overnight and treated with the indicated compounds for an additional 24 hours. Cells were trypsinized and resuspended in complete media in the presence of indicated compounds in tubes and the cell suspension was incubated for 60 min at 37 °C in a 5% CO_2_ incubator. Cells were then plated onto glass-bottom plates (Fisher Scientific, #NC0397150) coated with 50 μg/mL collagen-I (Millipore Sigma, #CLS354236) and allowed to spread for 60 min in the presence of the indicated compounds.

Following spreading, cells were fixed with 4 % paraformaldehyde in 1X PBS for 15 min at room temperature. Immunostaining was performed as previously described (44). Briefly, cells were permeabilized with 0.1 % Triton X-100 for 5 min, washed three times with 1X PBS, and blocked for 60 min at room temperature in a blocking buffer containing 5% normal goat serum and 2 % BSA in 1X PBS. Cells were incubated with anti-vinculin primary antibody (ProteinTech, #26520-1-AP; 1:200 dilution) for 60 min at room temperature, gently washed with 1X PBS three times, and then incubated for 45 min with Alexa Fluor-conjugated secondary antibodies (1:200 dilution) (Thermo Fisher Scientific, #A-11008) combined with Phalloidin (1:200 dilution) (Abcam, #ab176759). After three final washes, cells were mounted in ProLong Gold Antifade Mountant (Thermo Fisher Scientific, #P36984) and imaged using a Dragonfly 200 High-speed Spinning Disk confocal imaging platform (Andor Technology Ltd) mounted on a Leica DMi8 microscope stand, equipped with a 100X/1.4 oil immersion objective, an iXon EMCCD camera, and an sCMOS Zyla camera. Data acquisition and simultaneous deconvolution were performed using Fusion software (Version 2.3.0.36, Oxford Instruments) and Imaris, respectively. Cell area was quantified from maximum-intensity projected single-cell TIFF images by manually outlining each cell and measuring the enclosed area in Fiji/ImageJ. Focal adhesion numbers (FAs) were calculated from individual cells using vinculin as a focal adhesion marker. Automated FA detection was performed using a custom Fiji/ImageJ macro generated with the Record function. The workflow included following commands setAutoThreshold(“Yen dark 16-bit no-reset”); setThreshold(7279, 65535, “raw”); run(“Convert to Mask”); run(“Set Measurements…”, “area mean standard min add redirect=to the .tif decimal=3”); run(“Analyze Particles…”, “size=15-Infinity pixel show=Outlines display exclude overlay”). The final mask was further compared to the raw .tif image to visually confirm the results. The result gained from the automated analysis was also validated by manual counting.

## Acknowledgements

We thank ALBA for providing the synchrotron-radiation facility and the staff of the XALIC-13 beamline for their assistance in the data collection. DL has support from Worldwide Cancer Research (25–0168) and from a Generation of Knowledge Grant (PID2024-161627NB-I00), funded by MICIU/AEI/10.13039/501100011033 and by FEDER/UE.

This research was supported by the Cellular Imaging Shared Resource (RRID:SCR_022609) and Flow Cytometry Shared Resource (RRID:SCR_022613) of the Fred Hutch/University of Washington/Seattle Children’s Cancer Consortium (P30 CA015704).

This work was supported by the Robert L. Fine Cancer Research Foundation grant (B.N.). The content is solely the responsibility of the authors and does not necessarily represent the official views of the Robert L. Fine Cancer Research Foundation.

