## Supplementary Materials for "*De novo* design of selective kinase modulators"

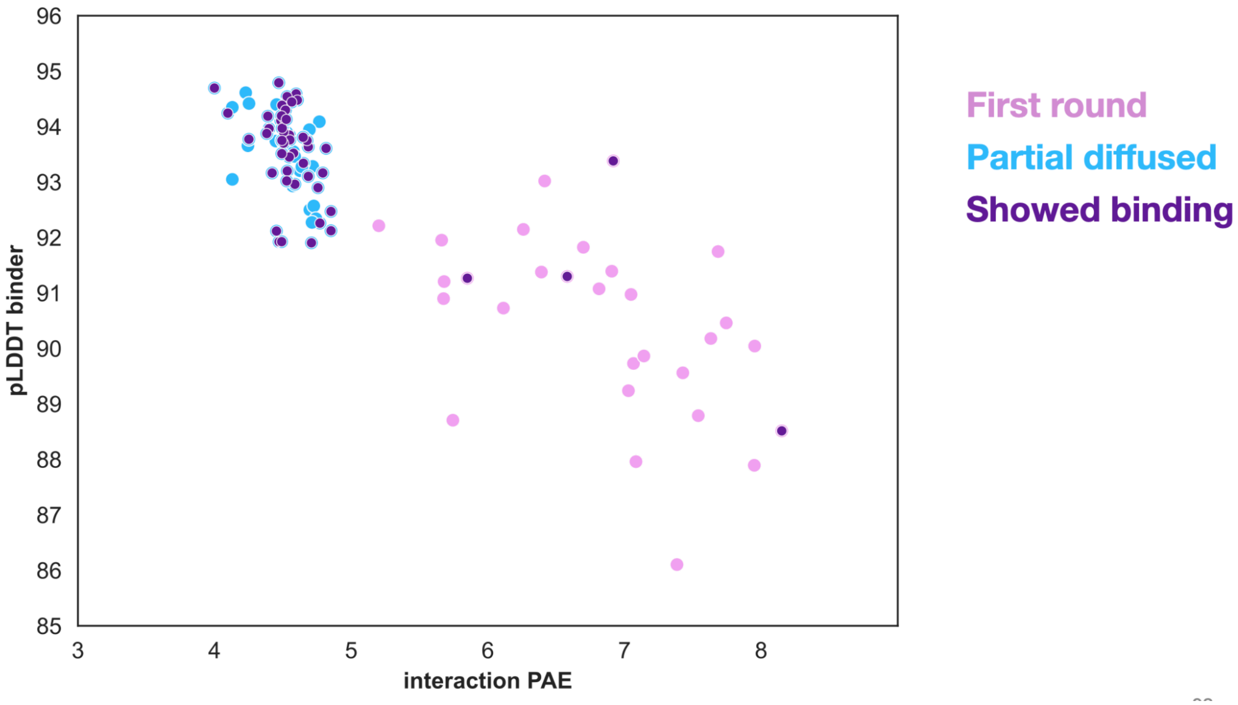

**Figure S1. Computational confidence metrics for designed kinase binders.** Binder pLDDT is plotted against interaction predicted aligned error (PAE) for designs from the first design round (pink) and partially diffused designs (cyan). Designs that showed experimental binding are highlighted in purple. Each point represents one design; higher pLDDT and lower interaction PAE indicate greater predicted structural and interaction confidence, respectively.

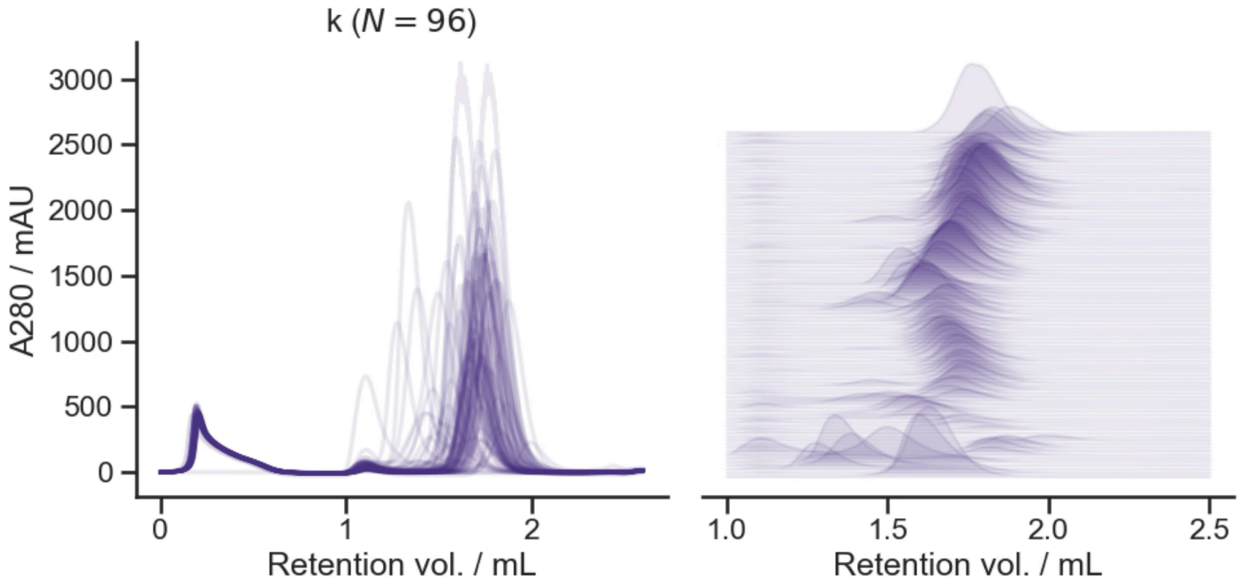

**Figure S2. Size-exclusion chromatography analysis of proteins produced using the Semi-Automated Protein Production (SAPP) workflow.** Analytical SEC profiles are shown for 96 protein-production samples. Overlaid absorbance traces measured at 280 nm are presented as conventional chromatograms (left) and as a vertically offset waterfall plot focused on the principal elution region (right). Each trace represents one sample.

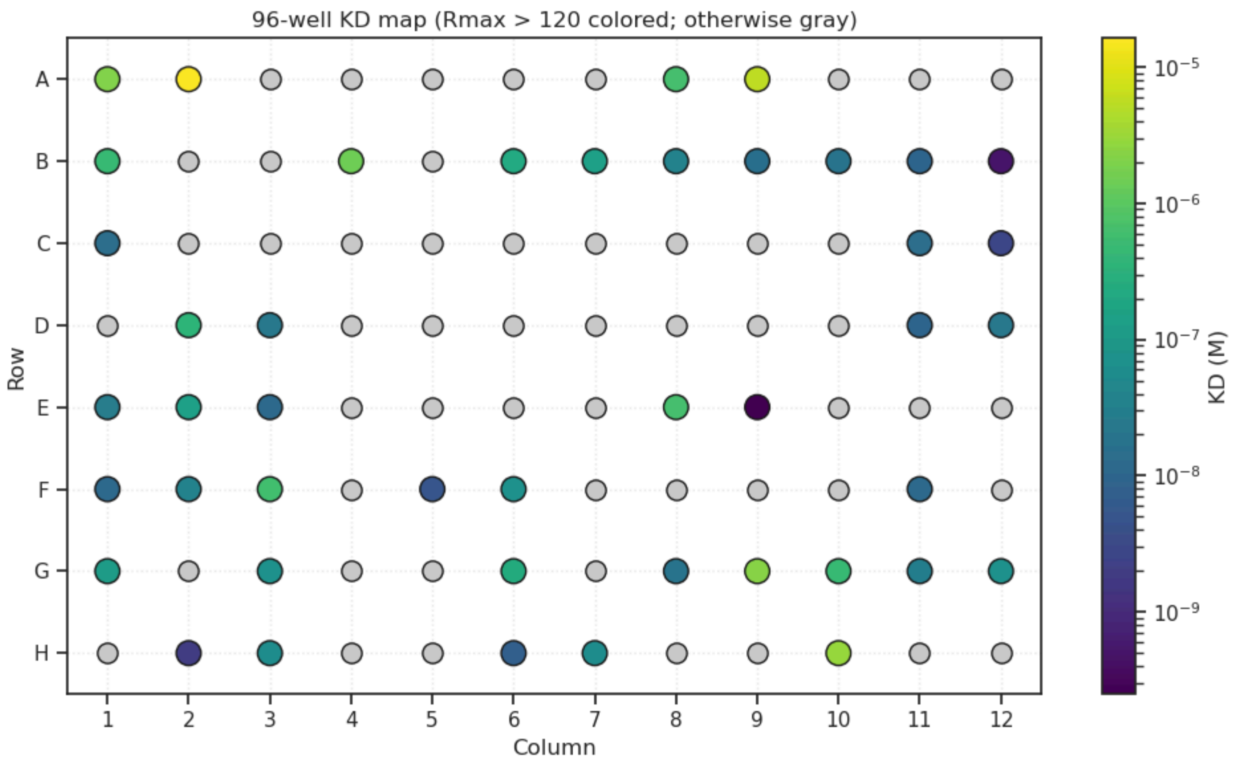

**Figure S3. Surface plasmon resonance affinity map for the 96-well screening plate.** Each circle represents one well. Wells with a fitted maximum response, $R_{max}$, greater than 120 are colored according to the fitted equilibrium dissociation constant, K_D, on a logarithmic molar scale; wells below this response threshold are shown in gray and were discarded for further analysis. Lower K_D values indicate tighter binding.

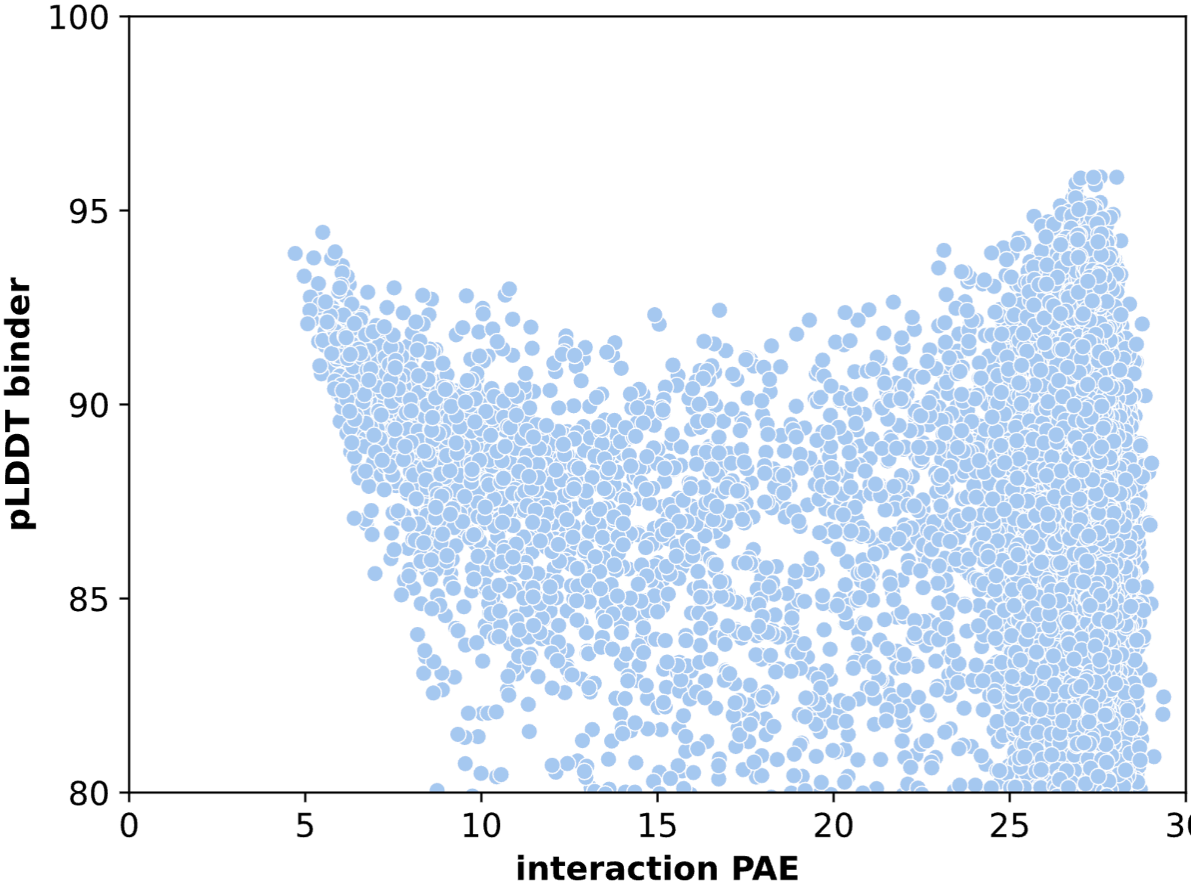

**Figure S4. Computational confidence landscape for Src-targeting binder designs.** Binder pLDDT is plotted against interaction PAE for the predicted Src-binder complexes. Each point represents one design. The distribution shows the range of binder-structure confidence and predicted interchain alignment error within the design pool; higher pLDDT and lower interaction PAE correspond to more confident predicted complexes.

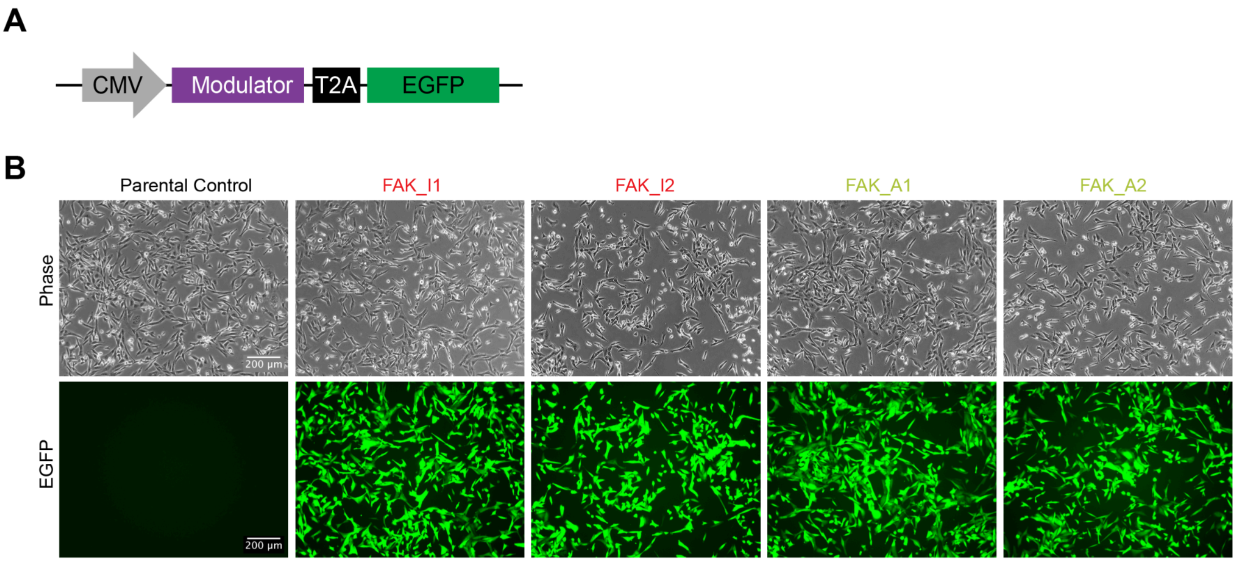

**Figure S5. Expression of *de novo* FAK modulators in MDA-MB-231 cells. (A)** Schematic of design of lentiviral plasmids to express *de novo* FAK modulators. EGFP is expressed from the same reading frame of the *de novo* FAK modulators coding sequence, separated by T2A self-cleaving peptide site. **(B)** Representative images of MDA-MB-231 parental control cells or MDA-MB-231 cells stably expressing FAK_I1, FAK_I2, FAK_A1 or FAK_A2.

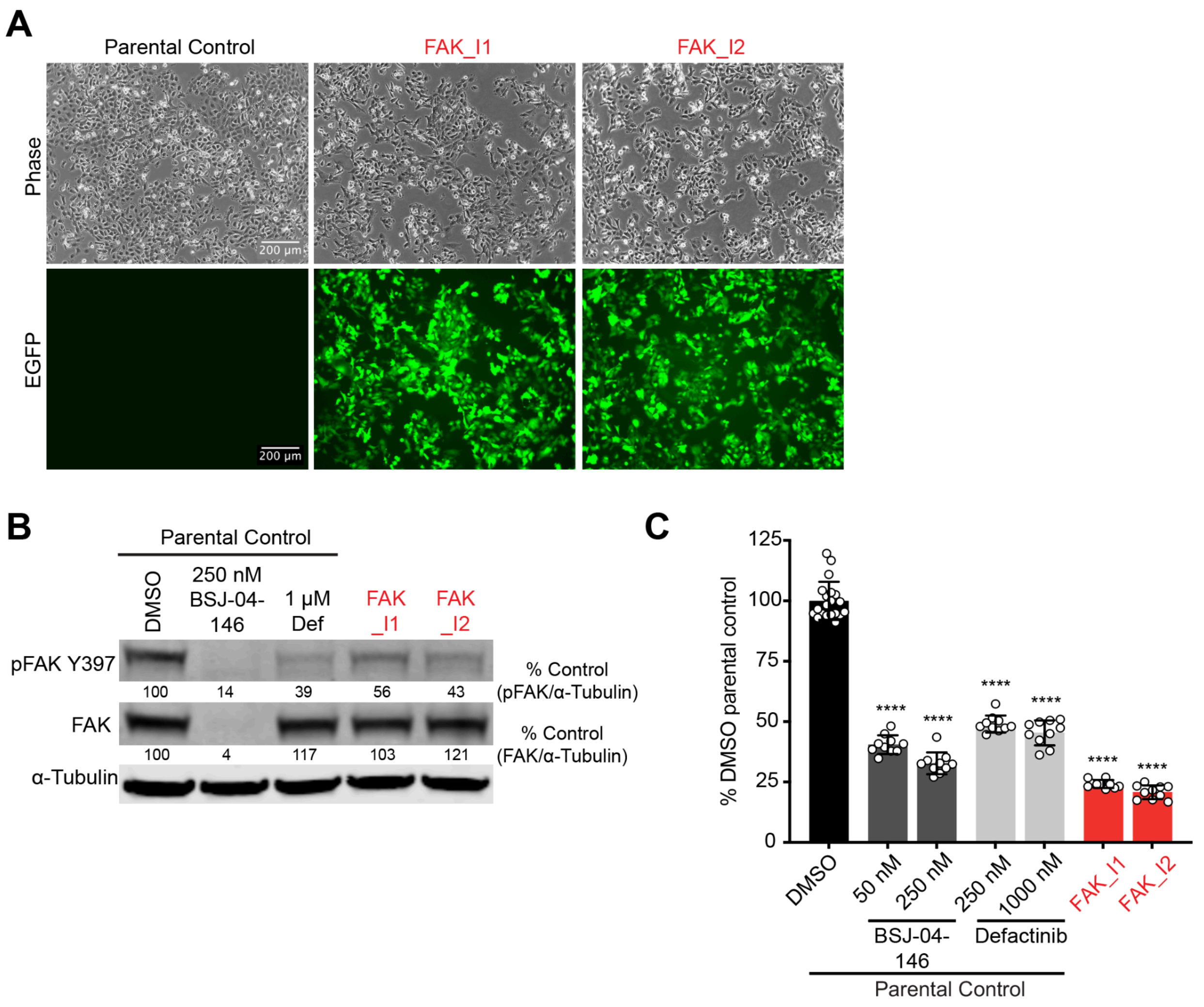

**Figure S6. Expression and characterization of *de novo* FAK modulators in PATU-8988T cells. (A)** Representative images of PATU-8988T parental control cells or PATU-8988T cells stably expressing FAK_I1, or FAK_I2. **(B)** Immunoblot analysis of the indicated PATU-8988T cells. PATU8988T parental control cells were treated with DMSO, 250 nM BSJ-04-146 or 1 mM defactinib for 24 hours. PATU8988T cells stably expressing FAK_I1 or FAK_I2 were treated with DMSO for 24 hours. **(C)** DMSO-normalized antiproliferation of the indicated PATU-8988T cells cultured as ultra-low adherent 3D spheroid suspensions for 120 hours. PATU-8988T parental control cells were treated with DMSO or the indicated doses of BSJ-04-146 or defactinib. PATU-8988T cells stably expressing FAK_I1 or FAK_I2 were treated with DMSO. Data are presented as the mean ± SD of 10-20 biologically independent samples and are representative of $n$ = 3 independent experiments. ****P < 0.0001 is indicated after analysis using 1-way ANOVA with post hoc Dunnett's test.

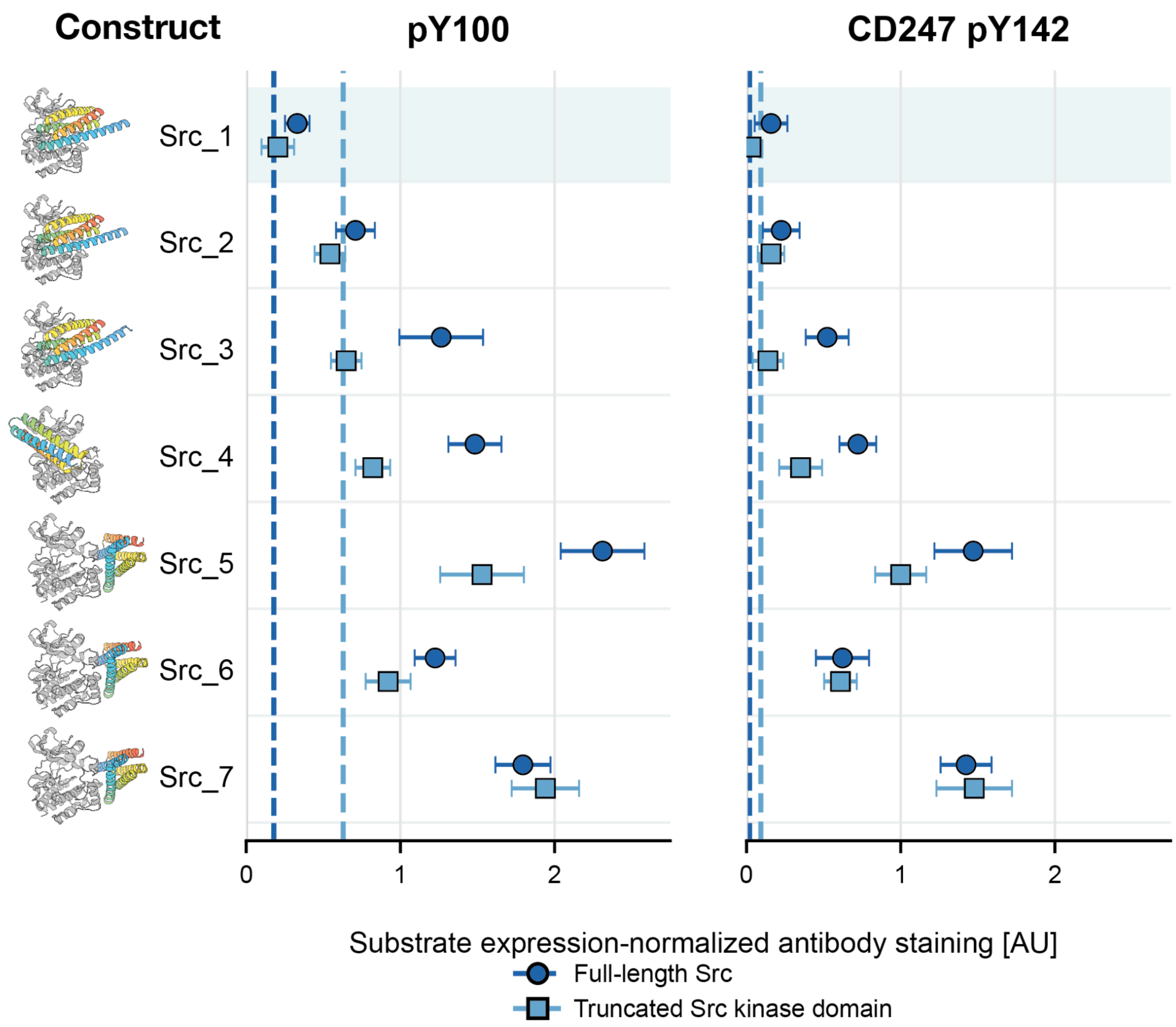

**Figure S7. Tethered Src-directed designs differentially suppress broad and reporter-specific tyrosine phosphorylation.** Seven Src-directed designs (Src_I1 - Src_I7) were tethered to either constitutively active full-length Src (FL) or a truncated Src kinase-domain construct (TRC) and co-expressed with a CD247 ITAM reporter. Left, predicted Src-design complex models aligned on the Src chain and shown in a common orientation. Middle, reporter-expression-normalized pY100 staining, providing a broad measure of cellular tyrosine phosphorylation. Right, reporter-expression-normalized CD247 pY142 staining, reporting phosphorylation of a defined site within the CD247 ITAM reporter. Dark-blue circles denote FL Src and light-blue squares denote TRC Src. Dark- and light-blue dashed vertical lines indicate the corresponding PPP2CA pseudosubstrate benchmark for the FL and TRC contexts, respectively, within each readout. Points at or to the left of the corresponding dashed line have signals comparable to or lower than the PPP2CA benchmark. Lower values indicate stronger suppression of Src-associated phosphorylation. Src_I1 produced the strongest and most consistent suppression, Src_I3 and Src_I4 showed greater suppression with TRC than with FL Src, and Src_I5 and Src_I7 showed little suppression.

| **Hotspot residue** |
|---|
| 596 |
| 473 |
| 440 |
| 516 |
| 576/577 |
| 649 |
| 657 |
| 423 |
| 584 |

**Supplementary Table T1. Residue numbering of hotspot positions used in this study.** All residue positions are reported according to the corresponding UniProt reference sequence.

| | ***FAK kinase+FAK_A1*** |
|---|---|
| **Data collection** | |
| X-ray source | ALBA, BL13-XALOC |
| Wavelength (Å)<br>Space group<br>Cell dimensions | 0.920241<br>$P22_12_1$ |
| a, b, c (Å)<br>a, b, g (°) | 42.22, 87,86, 109,94<br>90, 90, 90 |
| Resolution (Å) | 46.60-2.04 (2.10-2.04)* |

| | |
|---|---|
| Total reflections | 341160 (25222) |
| Multiplicity | 12.7 (12.4) |
| Unique reflections | 26844 (2038) |
| Completeness (%) | 99.9 (99.1) |
| $R_{merge}$ (%) | 8.0 (212.3) |
| $R_{meas}$ (%) | 8.3 (221.5) |
| $R_{pim}$ (%) | 2.4 (62.4) |
| CC(1/2) | 0.999 (0.899) |
| *I* / s*I* | 18.0 (1.6) |
| **Refinement** | |
| Resolution (Å) | 46.60-2.04 |
| Reflections (work/test set) | 25423/1339 |
| $R_{factor}$ / $R_{free}$ (%) | 24.0/26.6 |
| No. atoms | 2856 |
| Protein | 2784 |
| Solvent | 72 |
| Other | 0 |
| R.m.s. deviations | |
| Bond lengths (Å) | 0.003 |
| Bond angles (°) | 0.993 |
| mean *B* value (Å$^2$) | 50.48 |

**Supplementary Table T2.** Crystallographic data collection and refinement statistics. *Values in parentheses are for highest-resolution shell.
